# Human adipocyte differentiation and composition of disease-relevant lipids are regulated by miR-221-3p

**DOI:** 10.1101/2020.06.30.180083

**Authors:** Maria A. Ahonen, Muhammad Yasir Asghar, Suvi J. Parviainen, Gerhard Liebisch, Marcus Höring, Marjut Leidenius, Pamela Fischer-Posovszky, Martin Wabitsch, Tomi S. Mikkola, Kid Törnquist, Hanna Savolainen-Peltonen, P.A. Nidhina Haridas, Vesa M. Olkkonen

## Abstract

MicroRNA-221-3p (miR-221-3p) is associated with both metabolic diseases and cancers. However, its role in terminal adipocyte differentiation, adipocyte lipid metabolism, and lipid signaling in human disease are uncharacterized. miR-221-3p or its inhibitor were transfected into differentiating or mature human adipocytes. Triglyceride (TG) content and adipogenic gene expression were monitored, global lipidome analysis was carried out, and mechanisms underlying the effects of miR-221-3p were investigated. Finally, cross-talk between miR-221-3p expressing adipocytes and MCF-7 breast carcinoma (BC) cells was studied, and miR-221-3p expression in tumor-proximal adipose biopsies from BC patients analyzed. miR-221-3p overexpression inhibited terminal differentiation of adipocytes, as judged from reduced TG storage and gene expression of the adipogenic markers *SCD1, GLUT4, FAS, DGAT1/2, AP2, ATGL* and *AdipoQ*. The signaling adaptor protein 14-3-3γ was identified as a potential mediator of the miR-221-3p effects. Importantly, miR-221-3p overexpression inhibited *de novo* lipogenesis but increased the concentrations of ceramides and sphingomyelins, while reducing diacylglycerols, concomitant with suppression of sphingomyelin phosphodiesterase, ATP citrate lyase, and acid ceramidase. miR-221-3p expression was elevated in tumor proximal adipose tissue from patients with invasive BC. Conditioned medium of miR-221-3p overexpressing adipocytes stimulated the invasion and proliferation of BC cells, while medium of the BC cells enhanced miR-221-3p expression in adipocytes. miR-221-3p plays a pivotal role in the terminal differentiation of white adipocytes and their lipid composition. Elevated miR-221-3p impairs adipocyte lipid storage and differentiation, and modifies their ceramide, sphingomyelin, and diacylglycerol content. These alterations are relevant for metabolic diseases but may also affect cancer progression.

## Introduction

Adipose tissue (AT) is a *bona fide* endocrine organ, which regulates energy homeostasis of the body [1]. Defects in adipocyte lipid storage and metabolism contribute to the pathogenesis of many diseases including insulin resistance, non-alcoholic fatty liver disease (NAFLD), type 2 diabetes, and several forms of cancer [2–5]. Adipocyte differentiation is a complex process orchestrated by a series of signaling and transcriptional events. Adipocytes are initially derived from mesenchymal stem cells, which commit to form preadipocytes that further differentiate to lipid-laden, mature adipocytes [6, 7]. Mature adipocytes also have the potential to de-differentiate into pluripotent cells or other cell types [8, 9]. Dysregulation of adipogenesis or defects in adipocyte lipid storage lead to ectopic lipid accumulation resulting in insulin resistance, fatty liver disease, and eventually to type 2 diabetes. In obesity, adipocyte lipid storage capacity is exceeded and abundant lipids are routed to ectopic sites, while in the tumor microenvironment adipocytes are shown to delipidate and provide fatty acids to malignant cells for cancer progression [10–12]. Adipocyte lipid composition also plays a role in regulating the function of AT as well as in the progression of disorders such as NAFLD and type 2 diabetes [13, 14]. Recently a number of studies have shown that microRNAs (miRNAs) plays major roles in maintaining adipocyte physiology and contribute to the development of metabolic diseases [15–18].

MicroRNAs are short 18-25 nucleotide long non-coding RNAs, which alter protein expression via inducing mRNA degradation or inhibiting translation by physically binding to mRNA [19, 20]. A number of miRNA species are known to regulate adipogenesis, adipocyte lipid storage, metabolism, and adipokine secretion by directly targeting protein expression of key genes involved in signaling and transcriptional regulation. MicroRNA-221-3p (miR-221-3p) is one such miRNA shown to affect adipocyte differentiation, metabolism and insulin signaling [21–24]. miR-221 expression is elevated in obesity and it is induced upon AT inflammation [21, 23]. Furthermore, miR-221-3p is considered a cancer-associated miRNA, oncomiR, due to its abundant expression in cancerous tumors as well as in the circulation of patients suffering from several forms of cancer. Consequently, miR-221 and its family member sharing the same seed sequence, miR-222, are emerging as prognostic markers for various cancers [25–27]. Although miR-221-3p is known to affect adipocyte metabolism, the mechanisms by which it affects adipocyte differentiation are not known. Moreover, its effect on adipocyte lipid composition has not been investigated. Since miR-221-3p is a potential candidate to identify hitherto unknown mechanisms of adipogenesis, adipocyte lipid composition, and their distortions in human diseases, we assessed in this study the effects of miR-221-3p on human adipocyte differentiation and lipid composition, as well as the underlying mechanisms.

## Materials and methods

### Subjects and study design

Subcutaneous adipose tissue biopsies were obtained from women undergoing nonmalignant breast surgery (reduction mammoplasty; n=30). To profile miR-221-3p and adipocyte marker gene expression in breast adipose tissue proximal to tumors (distance to the tumor < 5cm), biopsies were withdrawn from women operated for breast cancer (n=47). The subjects’ height, weight, waist and hip circumferences, medical history and use of medications were recorded at the preoperative visit (Table 1). AT biopsies of 1 g were obtained during the operation from the reduction mammoplasty or mastectomy patients. The biopsies were taken proximally to the breast tumor from the mastectomy patients. The samples were snap frozen in liquid nitrogen and stored at −80°C. A written informed consent was acquired from all study subjects. The study was approved by the Ethics Committee of Helsinki University Central Hospital.

**Table 1.** Clinical characteristics of subjects undergoing reduction mammoplasty or mastectomy.

|  | Reduction mammoplasty<br>(n=30) | Breast cancer (n=47) |
| --- | --- | --- |
| Age, y<sup>a</sup> | 46 (21-77) | 65 (29-89) |
| Postmenopausal, %<sup>b</sup> | 45 (13) | 68 (32) |
| BMI, kg/m<sup>2a</sup> | 28 (18-38) | 25 (19-40) |
| WHR A.U. <sup>a</sup> | 0.88 (0.70-0.99) | 0.87 (0.63-0.99) |
| Hypertension, %<sup>b</sup> | 17 (5) | 30 (14) |
| Hypercholesterolemia, %<sup>b</sup> | 14 (4) | 23 (11) |
| Type 2 diabetes, %<sup>b</sup> | 7 (2) | 13 (6) |

<sup>a</sup> The data are expressed as median (range).
<sup>b</sup> The data are expressed as percentages (n).

### Cell Culture, transfections and collection of conditioned media

The roles of miR-221-3p and 14-3-3γ (encoded by the gene *YWHAG*) in adipogenesis were studied by transfecting adipocytes with miR-221 mimic or by silencing *YWHAG*. Simpson-Golabi-Behmel syndrome (SGBS) preadipocytes were cultured and transfected on day 6 of differentiation, followed by differentiation until day 13 or 14 according to Fischer-Posovszky et al. [28]. Transfections were carried out with 100 nM miRNA-221-3p mimic (Qiagen MSY0000278), anti-miR-221-3p (Qiagen MIN0000278) or non-targeting siRNA (NT; Qiagen 1027280), or alternatively, with 100 nM *YWHAG* siRNA (Qiagen SI00100653) or Silencer Select^™^ non-targeting control 2 (SS2; Thermo Fisher Scientific 4390846, Waltham, MA). Transfections were performed by using the RNAiMax^™^ reagent (Thermo Fisher Scientific). Transfection complexes were incubated on cells for 48 hrs, and the differentiation was continued until mature adipocytes (day 14-16) were lysed or fixed for further experiments.

To study the effect of miR-221-3p on metabolism in mature adipocyte, SGBS preadipocytes were differentiated for 13 to 14 days as specified above and transfected with 200 nM miR-221-3p mimic or non-targeting control for 72 hrs. Cells were then used for measurement of [^3^H]acetic acid incorporation into lipids or lysed for gene and protein expression or lipidomic analyses. For generating conditioned medium from miR-221-3p transfected mature adipocytes, cells were carefully washed after 48 hrs of transfection, and transfection complexes were replaced with serum free DMEM F-12 (Gibco 31330-038). The freshly added medium was incubated on the cells for 24 hrs. Adipocyte conditioned media (ACM) was collected and centrifuged at 300 x g for 5 min, at 1500 x g for 10 min and stored at -20 °C until further use for experiments.

SGBS adipocytes were treated with breast cancer cell conditioned medium (CCM). Michigan Cancer Foundation 7 (MCF-7) cells were cultured with DMEM (Sigma D6429) with 10% foetal bovine serum (FBS), 2% L-glutamine, penicillin/streptomycin. Cells were cultured to >80% confluency and used for obtaining conditioned medium. For obtaining the CCM, serum free DMEM (SFM; Sigma D5546) was added on MCF-7 cells for 24 hrs, followed by centrifugation as described above for ACM. SGBS cells were differentiated and, once matured, washed carefully. MCF-7 CCM or SFM were added on the adipocytes for 18 hrs and the cells were harvested for following experiments.

### Invasion assay

Invasion assays were performed on 6.5 mm diameter Transwell^™^ inserts (Corning Costar, Corning, NY) with pore size of 8 μm. The membranes were coated with 5μg/cm^2^ human collagen IV and allowed to dry overnight. They were then reconstituted with SFM at 37^°^C for 1 hr prior to the experiment. 50,000 MCF-7 cells were allowed to migrate/invade in 40% adipocyte conditioned medium (ACM) in SFM towards 60% normal medium containing 10% FBS and 40% of ACM for 24 h. Mitomycin C, (0.5 mg/ml; Sigma, M4287) was added in both lower and upper chambers to block proliferation. Next, the cells on top of the membrane were wiped off with a cotton swab. The migrated cells were fixed in 2% paraformaldehyde for 10 min and stained with 0.1% crystal violet in 20% methanol for 5 min. The membranes were rinsed with phosphate-buffered saline (PBS) and water and allowed to dry overnight. The cells were counted at 40x magnification in eight microscopic fields in a straight line bisecting the membrane.

### Proliferation assay

Proliferation of MCF-7 cells after ACM incubation (as explained above) was quantified by measuring [^3^H]thymidine incorporation. 50,000 cells were seeded on 35-mm plates and allowed to grow for overnight. Next day, the medium was changed to 40% ACM in culturing medium Four hours prior to the end of each experiment, 0.4 μCi/ml [^3^H]thymidine was added to each culture plate. The cells were washed three times with PBS, incubated for 10 min with 5 % trichloric acetic acid, and then incubated for 10 min with 0.1 m NaOH. The samples were transferred into scintillation tubes and high sample load scintillation cocktail Optiphase Hisafe 3 was added. The radioactivity was measured using a Wallac 1414 liquid scintillation counter.

### Triglyceride measurement and lipid droplet staining

TG content was measured with an enzymatic GPO-PAP assay kit (Cobas, Roche/Hitachi, Tokyo, Japan) according to the manufacturer’s protocol. TG levels were normalized to protein content. Transfected and differentiated SGBS adipocytes were fixed on coverslips with 4% paraformaldehyde in PBS, followed by washing with cold PBS. The coverslips were stained with Bodipy 493/503 (Molecular Probes/Thermo Scientific, Eugene, OR) at room temperature for 20 minutes and mounted with Moviol (Calbiochem, La Jolla, CA) containing 5 μg/ml DAPI (Thermo Scientific/Molecular Probes) and 50 mg/ml 1,4-Diazabicyclo-[2.2.2] octane (Sigma-Aldrich). Coverslips were scanned with 3D Histech Pannoramic scanner (3DHISTECH, Budapest, Hungary) and 20x snapshots taken using CaseViewer (version 2.2; 3DHISTECH). Approximately 500 cells were analyzed by using Fiji software and MRI_Lipid Droplets tool macro (http://dev.mri.cnrs.fr/projects/imagej-macros/wiki/Lipid_Droplets_Tool).

### Gene and miRNA expression analyses

Gene and miR-221-3p expressions were investigated by using quantitative real time PCR (qPCR). Total RNA from SGBS cells or from tissue biopsies was isolated with Qiagen (Gaithersburg, MD) kits according to the manufacturer’s protocols. SuperScript^®^ VILO™ synthesis Kit (Invitrogen, Carlsbad, CA) was used for reverse transcription of cDNA for gene expression analysis. To detect mRNA expression, qPCR was conducted using Roche SYBR-Green^®^ master mix and a LightCycler 480 II Real-Time PCR system (Roche Applied Science, Penzberg, Germany). For analysis, crossing point (Cp) values were calculated from amplification curves and normalized to Cp values of the housekeeping genes succinate dehydrogenase complex subunit A (*SDHA*) and actin. Primer sequences are specified in Table 2. To measure expression of miR-221-3p, RNA was reverse transcribed by using TaqMan^®^ miRNA reverse transcription kit (Applied Biosystems, Foster City, CA) and qPCR was carried out with TaqMan^®^ microRNA assay according to the manufacturer’s protocol, using miR-221-3p assay (000524), and RNU44 assay (001094) as a house keeping control.

**Table 2.**
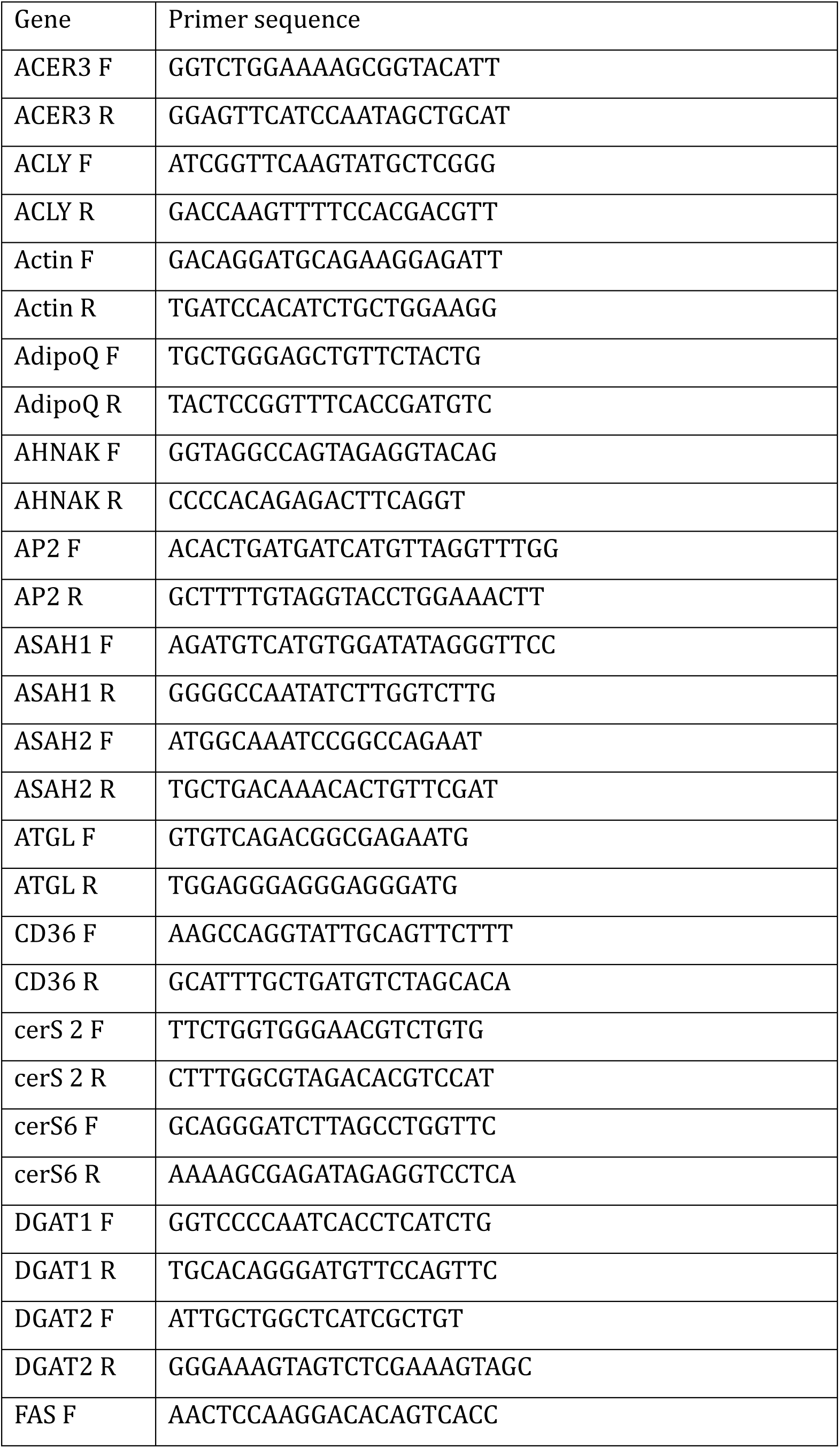
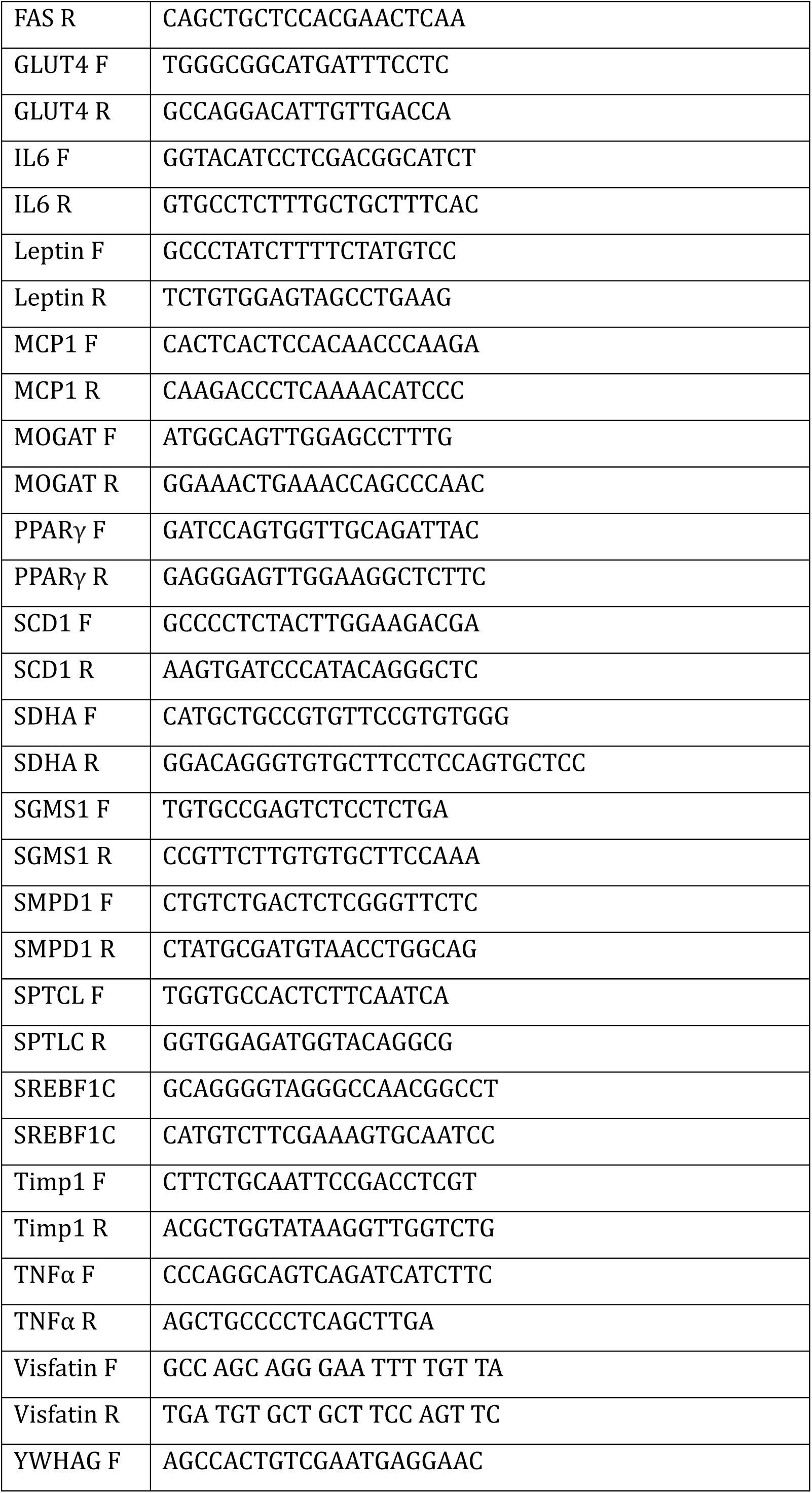
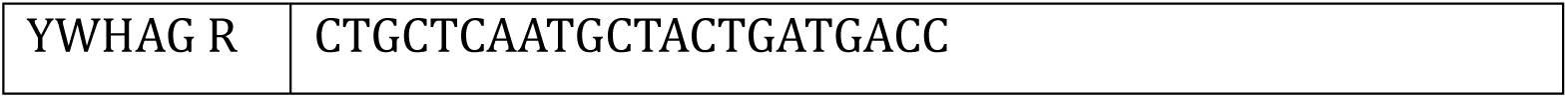
Sequences of primers used for qPCR analysis.

### Western blotting

SGBS protein expression was quantified by western blotting. Cells were lysed in RIPA buffer (15 mM Tris-HCl, pH 7.4, 1% NP40 1%, 1.25% sodium deoxycholate 1.25%, 150 mM NaCl, 1 mM EDTA, 1% SDS). Equal amounts of protein were loaded on 10% or 4-15% sodium dodecyl sulfate polyacrylamide gels (Fast Cast TGX Stain-Free or 4-15% Mini-Protean TGX Stain-Free gels, BioRad, Hercules, CA). Blotting was done on PVDF membranes using a BioRad Transblot system. Membranes were blocked and probed overnight with anti-SCD1 (Cell Signaling 2438S), anti-AdipoQ (Sigma A6354), anti-14-3-3γ (Proteintech 12381-1-AP), anti-ACLY (Santa Cruz Biotechnology SC-517267) or anti-ASAH1 (Santa Cruz Biotechnology SC-136275) in 5% milk in TBS with 0.5% Tween-20. Proteins were detected with enhanced chemiluminescence (ECL; Thermo Scientific, Waltham, MA). Image Lab (BioRad) was used to quantify the corresponding protein band intensities normalized to total protein intensity.

### Lipidome analysis

Lipids were extracted using Bligh and Dyer method [29]. Internal standards used were PC 28:0, SM 30:1,Cer d18:1/17:0, PE 40:0, DG 28:0, FC7D, CE 17:0, TG 51:0, LPC 13:0, LPE 13:0, Cer 35:1;2, HexCer 35:1;2, PC O-28:0, PG 28:0, PI 34:0, SM 30:1;2, CE 17:0 and PS 40:0. 10 mM ammonium acetate plus methanol/chloroform (3:1, v/v) (for low mass resolution tandem mass spectrometry), or chloroform/methanol/2-propanol (1:2:4 v/v/v) with 7.5 mM ammonium formate (for high resolution mass spectrometry) was used for solving the residues. Lipid analysis was conducted by direct flow injection analysis (FIA) and a hybrid quadrupole-Orbitrap mass spectrometer (FIA-FTMS; high mass resolution) [30] for diglycerides (DG) and TG or a triple quadrupole mass spectrometer (FIA-MS/MS; QQQ triple quadrupole) [31, 32] for the other classes. Lipid species were annotated according to the shorthand notation described previously [33]. Data was processed by using macros as previously described [34].

### [^3^H]Acetic acid labeling of lipids

SGBS adipocytes were differentiated and transfected on 6-well plates as specified above. The cells were then incubated with [^3^H]acetic acid (1 μCi/well; Amersham, GE Healthcare) in serum free DMEM F-12 for 4 hrs, after which the cells were washed and harvested into 2 % NaCl. The Bligh and Dyer method was used to extract the lipids [29], followed by thin layer chromatography by using hexane/diethyl ether/acetic acid/ water (70:45:1:0.25) as the running solution. Triolein, diolein and cholesterol were used as standards for TG, DG and cholesterol identification. The corresponding spots were scraped and radioactivity was determined by liquid scintillation counting.

### Statistical methods

Shapiro Wilk’s test was used to assess normality of the data. Correlations in the patient materials were studied by Spearman correlation test. The lipidomics data were studied using multiple t-test and significance determined by Bonferroni-Dunn method. The other *in vitro* data were analyzed using the Mann-Whitney U or Kruskal Wallis tests for non-normally distributed data and Student’s t-test or One-way ANOVA for normally distributed data. P-values ≤ 0.05 were considered significant and standard deviation was reported as standard deviation (SD) from the mean. Statistical analyses were conducted by using GraphPad Prism 8 (GraphPad Software, Inc., La Jolla, CA) or SPSS 25 (IBM SPSS Statistics for Macintosh, Version 25.0. Armonk, NY).

## Results

### miR-221-3p induces a differentiation defect in human adipocytes

miR-221-3p is reported to affect adipocyte differentiation when overexpressed at the initial stages of the process [24]. However, neither the mechanism by which it affects adipocyte differentiation, nor its impact on the later stages of the differentiation, has been investigated. Hence, we were interested in the effect of miR-221-3p on terminal adipocyte differentiation and the mechanism involved. We transfected miR-221-3p, anti-miR-221-3p or non-targeting (NT) control siRNA to SGBS adipocytes on day 6 of the differentiation and continued the differentiation until day 13 or 14 (Fig. 1A). A drastic, 42% reduction in the adipocyte TG content was observed upon miR-221-3p overexpression, while anti-miR-221-3p increased the TG content by 44% in comparison to NT (Fig. 1B), confirming that the endogenous miR-221-3p plays an important role in adipocyte TG storage. Consistently, when compared to the NT controls, the area of imaged lipid droplets per cell was reduced by 64% upon miR-221-3p transfection, whereas transfection with anti-miR-221-3p induced an increasing trend (+42%) in the lipid droplet area (Fig. 1C, D). We next wanted to confirm that miR-221-3p causes a differentiation defect, by conducting gene and protein expression analyses. Consistent with the phenotypic change, the mRNA expression of adipocytic differentiation marker genes (stearoyl-CoA desaturase-1 - *SCD1*; glucose transporter 4 - *GLUT4*; fatty acid synthase – *FAS*; diacyl glycerol acyl transferase1/2 - *DGAT1/2*; adipocyte fatty acid binding protein - *AP2*; adipocyte triglyceride lipase – *ATGL*; adiponectin – *AdipoQ*) were significantly decreased in cells transfected with miR-221-3p mimic. Moreover, expression of monoacylglycerol O-acyltransferase (*MOGAT*) and sterol-regulatory-element-binding protein 1c (*SREBF1C*) mRNA showed a reducing tendency but did not reach statistical significance (Fig. 2A). To confirm that the impacts of miR-221-3p are present at the level of the corresponding proteins, SCD1 and AdipoQ were analyzed by western blotting; both were significantly downregulated upon miR-221-3p mimic transfection (Fig 2B).

**Fig. 1.**
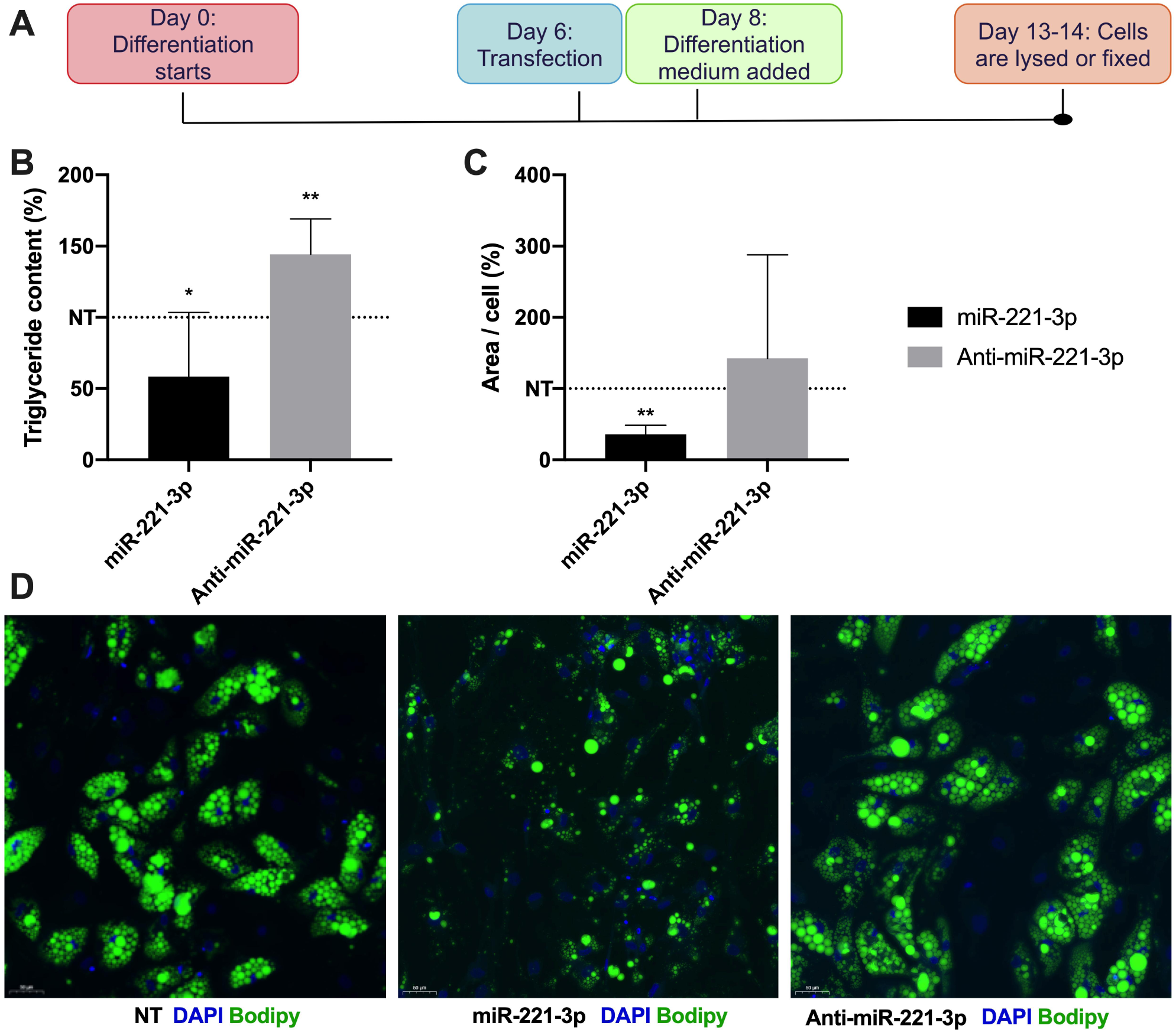
miR-221-3p attenuates lipid accumulation during SGBS adipocyte differentiation. A) To assess miR-221-3p’s role in terminal differentiation, adipocytes where transfected with non-targeting siRNA (NT), miR-221-3p mimic (miR-221-3p) or miR-221-3p inhibitor (anti-miR-221-3p) on day 6 of differentiation. B) Triglyceride content in SGBS on day 13 transfected on day 6 (n=8, two independent experiments) C) Average lipid droplet area per cell quantified from lipid droplet staining. Approximately 500 cells were quantified. D) Lipid droplet staining (Bodipy) of mature SGBS adipocytes transfected on day 6 (n=2, two independent experiments). The quantifications are expressed as percentages, where NT is 100% (indicated by a dotted line). The data is represented as mean with SD. Statistical significance is designated as *\*P ≤0*.*05, **P < 0*.*01*.

**Fig. 2.**
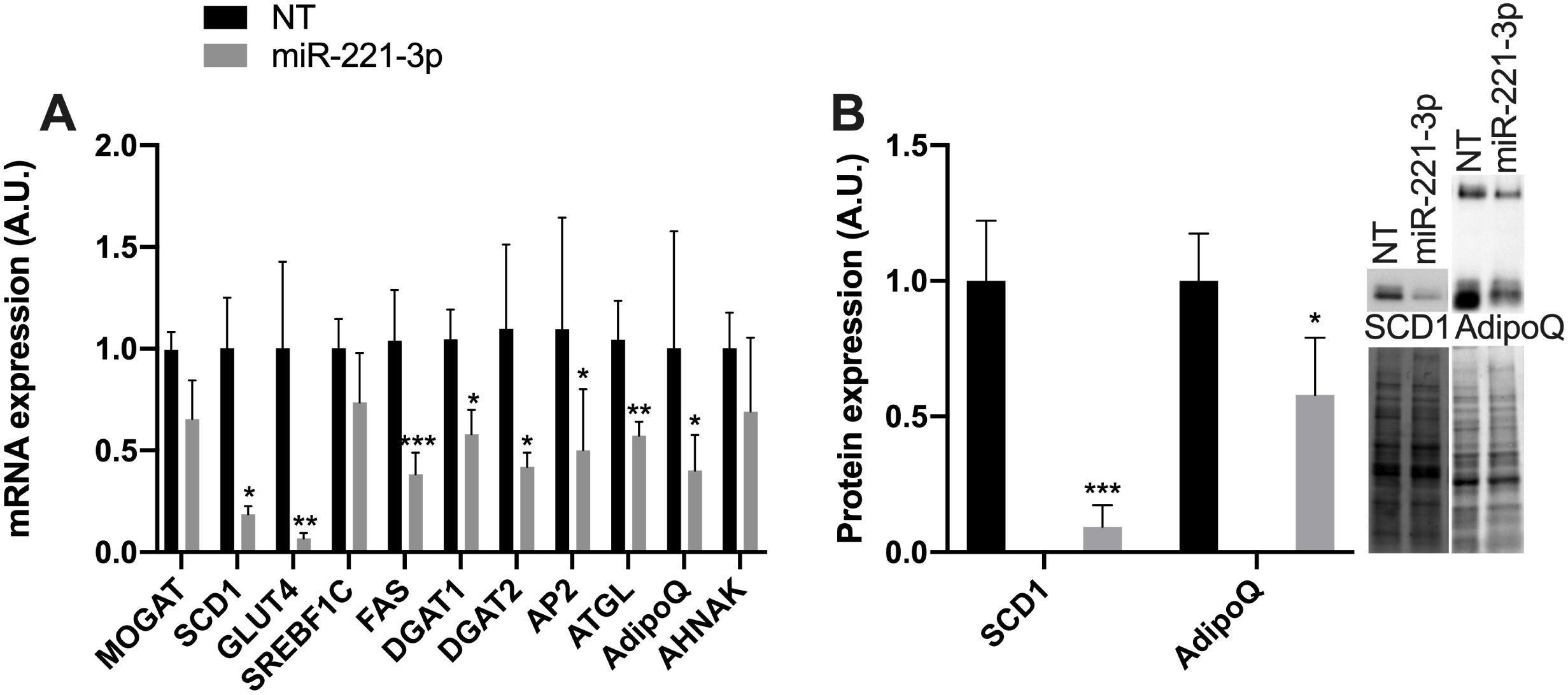
miR-221-3p dampens the terminal differentiation of adipocytes. A) mRNA expression of adipogenesis markers on day 13, after transfection of the adipocytes on day 6 of differentiation (n=5-7, two independent experiments). B) Quantification of protein expression of SCD1 and adiponectin (AdipoQ) in cells transfected with NT or miR-221-3p (n=7-8, two independent experiments). Representative western blots are shown on the right. Immunodetected bands are shown at the top and total protein below. The data is represented as mean with SD. Statistical significance is designated as *\*P ≤ 0*.*05, **P < 0*.*01, ***P < 0*.*001*.

### Adipocyte differentiation defect upon miR-221-3p overexpression may be mediated by down-regulation of 14-3-3γ

We next wanted to investigate how miR-221-3p dampens adipocyte differentiation. Several 14-3-3 proteins are connected to differentiation of various cell types, and 14-3-3ζ has been shown to regulate the adipogenic program [35]. We therefore hypothesized that also other 14-3-3 isoforms could play a role in adipocyte differentiation. miR-222, which shares an identical seed sequence with miR-221-3p (Fig. 3A), directly targets the *YWHAG* (encoding 14-3-3γ) 3’UTR [36], and also miR-221-3p is predicted (by miRWalk 2.0) to directly target 14-3-3γ [37]. Upon miR-221-3p transfection during SGBS adipocyte differentiation, a significant reduction of 14-3-3γ protein was observed, whereas antimiR-221-3p transfection resulted in an increased expression of 14-3-3γ (Fig 3B). We next knocked down 14-3-3γ to investigate whether this would mediate similar a phenotypic changes as miR-221-3p overexpression does. Indeed, the TG content of SGBS adipocytes was reduced by 25% and the lipid droplet area per cell by 46% upon depletion of 14-3-3γ (Fig. 3C). Inhibition of the adipogenic differentiation was also reflected in the mRNA markers of adipogenesis, *SCD1, GLUT4, FAS* and *AHNAK*, which were significantly downregulated upon knockdown of *YWHAG*/14-3-3γ (Fig. 4A). Since *YWHAG*/14-3-3γ has not been widely studied in human subcutaneous adipose tissue, we wanted to study whether *YWHAG* expression correlated with adipogenic markers in human white adipose tissue *in vivo*. This question was addressed by qPCR analysis of subcutaneous (breast) adipose tissue biopsies from 30 female human subjects. Significant positive correlation of the *YWHAG* mRNA with *FAS, SCD1* and *AHNAK* was observed (Fig. 4B, C, E).

**Fig. 3.**
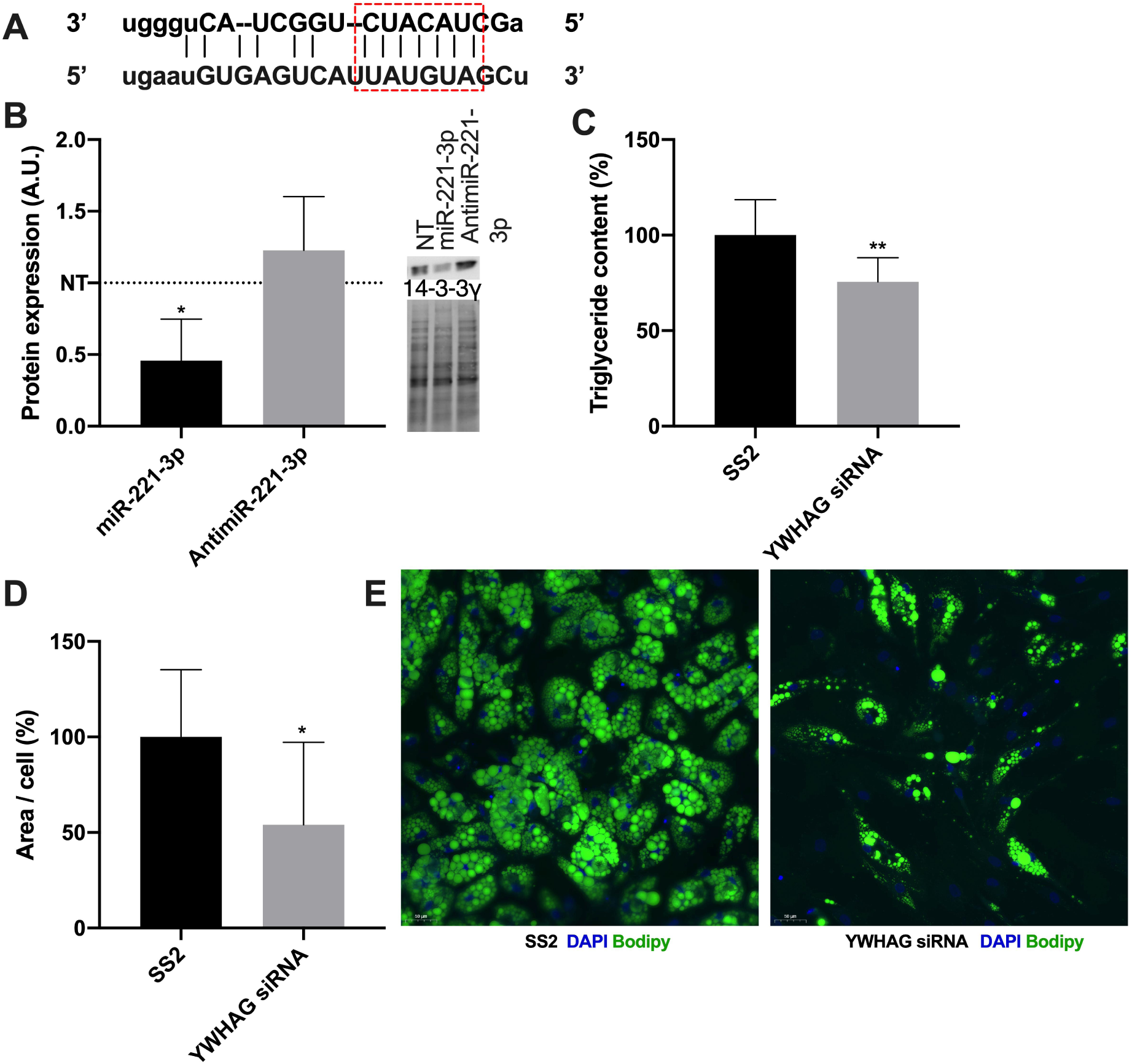
14-3-3γ, inhibited by miR-221-3p, affects lipid accumulation during the terminal differentiation of SGBS adipocytes. A) The seed sequence of miR-221-3p matches the 3’UTR of *YWHAG*, the mRNA encoding 14-3-3γ. B) Quantification of 14-3-3γ protein expression in mature SGBS transfected with non-targeting siRNA (NT), miR-221-3p mimic (mir-221-3p) or mir-221-3p inhibitor (anti-miR-221-3p; n=8 two independent experiments). The level in NT is indicated with a dotted dine. Representative immunoblots are shown on the right: immunodetected bands are displayed at the top and the total proteins in the lanes below. C) Triglyceride content in mature SGBS adipocytes transfected with Silencer Select 2^™^ non-targeting control (SS2) or *YWHAG* siRNA on day 6 of differentiation (n=9-10, two independent experiments). D) Lipid droplet area per cell quantified from the lipid droplet staining. Approximately 500 cells were quantified. E) Lipid droplet staining (Bodipy) of mature SGBS adipocytes transfected on day 6 of differentiation (n=2, two independent experiments). The data is represented as mean with SD. Statistical significance is designated as *\*P ≤ 0*.*05, **P < 0*.*01*.

**Fig. 4.**
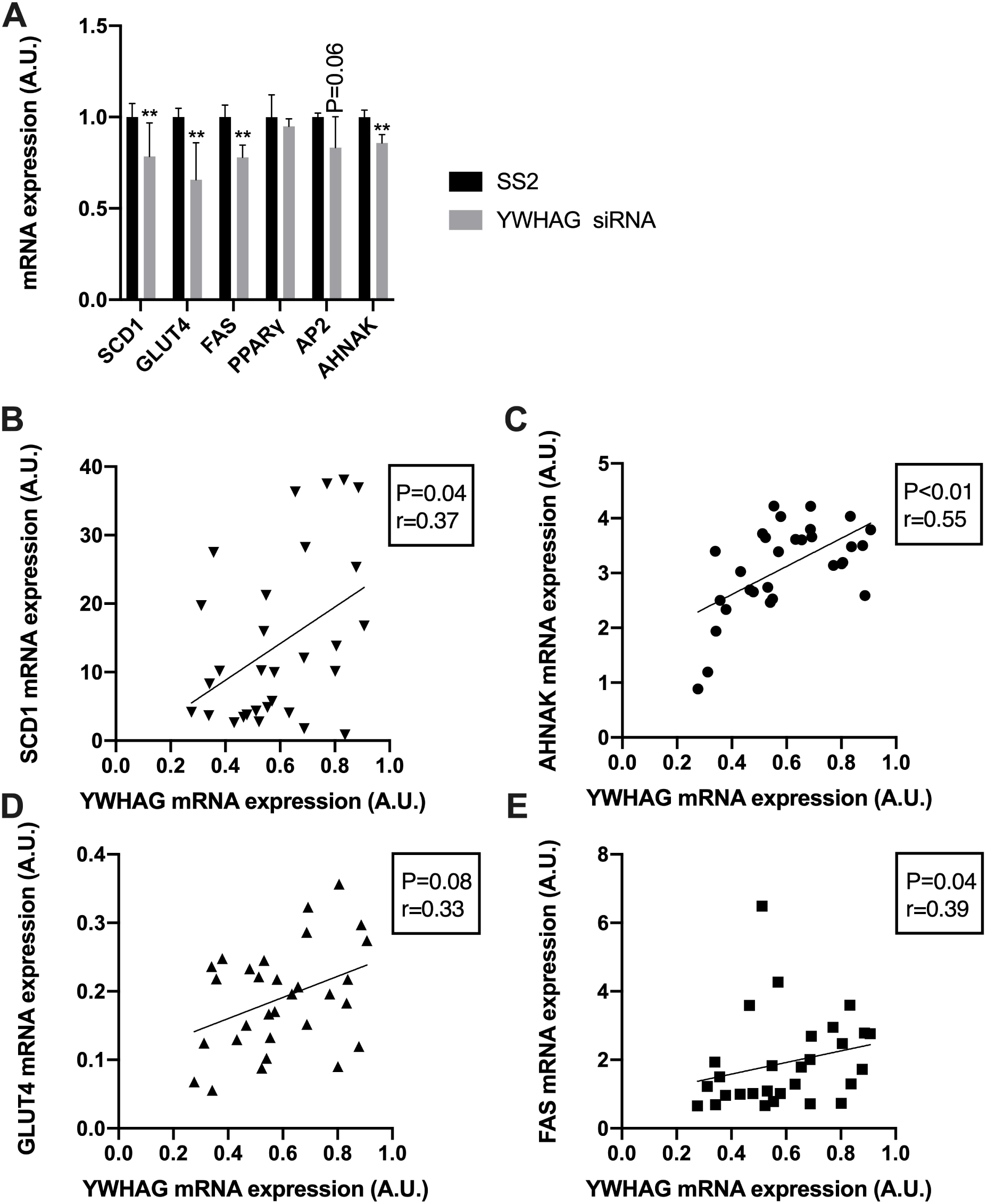
Association of the *YWHAG* mRNA (encoding 14-3-3γ) with adipocyte differentiation markers *in vitro* and *in vivo*. A) mRNA expression of adipocyte differentiation markers in mature SGBS cells that were transfected with Silencer Select 2^™^ non-targeting control (SS2) or *YWHAG* siRNA on day 6 of differentiation (n=6, two independent experiments). The data is represented as mean with SD. Statistical significance is designated as *\*\*P < 0*.*01*. B-E) *YWHAG* mRNA correlates with stearoyl-CoA desaturase-1 (*SCD1*; B), *AHNAK* protein (C), glucose transporter 4 (*GLUT4*; D) and fatty acid synthase (*FAS*; E) mRNA in human subcutaneous adipose tissue (n=30). The P and r values are indicated in the respective panels.

### miR-221-3p alters the composition of signaling lipids in human adipocytes

Since miR-221-3p is connected to adipose tissue dysfunction, which is characterized by an altered lipid composition and a distortion of the adipocytes’ differentiation status [16, 24, 38–40], we wanted to elucidate the effects of miR-221-3p overexpression (72-hr transfection; Fig. 5A) on the lipidome of mature SGBS adipocytes and on mRNA expression of adipocytic differentiation markers in these cells. The adipogenic mRNAs *SCD1, GLUT4, FAS, DGAT1*, -*2 and ATGL* as well as the anti-inflammatory adipokine *AdipoQ* were all significantly downregulated, whereas the pro-inflammatory cytokines (visfatin, interleukin *6 - IL6* and monocyte chemoattractant protein *1 - MCP1*) mRNAs were significantly induced (Fig. 5B). Consistent with the mRNA observation, the AdipoQ protein was significantly reduced (Fig. 5C). However, the TG content of the mature adipocytes showed a modest reduction, which did not reach significance (Fig. 5D).

**Fig. 5.**
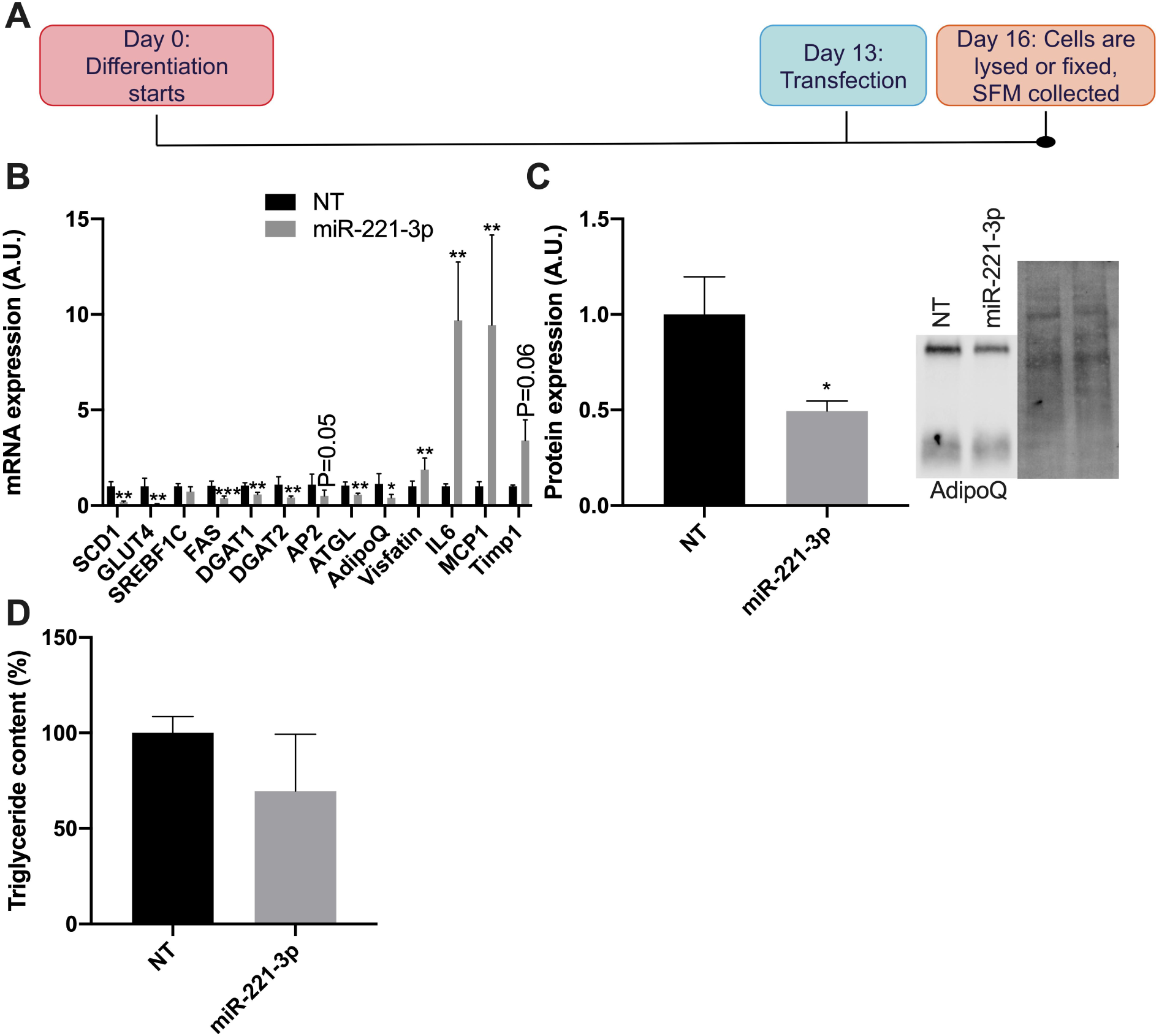
Impact of miR-221-3p on differentiation markers in mature adipocytes. A) SGBS adipocytes were transfected with non-targeting siRNA (NT) or miR-221-3p mimic (miR-221-3p) at the mature stage, on day 13 of differentiation. B) mRNA expression of adipocyte differentiation markers on day 16, after transfection on day 13 (n=4-7. 3 independent experiments). C) Quantification of protein expression of adiponectin (AdipoQ) in cells transfected with non-targeting siRNA (NT) or miR-221-3p mimic (miR-221-3p; n=4, two independent experiments). Representative western blots are shown: immunodetected bands on the left and total protein on the right. D) Triglyceride content in SGBS on day 16, after transfection on day 13 (n=6, two independent experiments). The data is represented as mean with SD. Statistical significance is designated as *\*P ≤ 0*.*05, **P < 0*.*01, ***P < 0*.*001*.

We next carried out a lipidomic analysis of miR-221-3p or NT transfected mature adipocytes. The miR-221-3p overexpressing adipocytes exhibited a significant reduction in the total diacylglycerol (DG) content and an accumulation of ceramides (Cer) and sphingomyelins (SM; Fig. 6A). Significantly affected Cer species were d18:1/22:0, d18:1/24:1, d18:1/20:0, d18:1/24:0, d18:1/18:0, d18:1/26:1; SM species 36:1, 40:1, 40:2, 38:2, 42:2, 42:1, 34:0, 36:2, and DG species 32:2, 34:2, 30:1, 32:1, 32:0, 34:1 (Fig. 6B). Of note, elevation of ceramides in white adipose tissue has previously been associated with obesity, insulin resistance and chronic inflammation in metabolic disease [13, 39–42], but also in a tumor environment [43]. Among the 183 glycerophospholipid species detected, only seven differed significantly between the miR-221-3p transfected and control adipocytes (data not shown), consistent with the view that the functional effect of miR-221-3p is quite specific for the sphingolipids and DG.

**Fig. 6.**
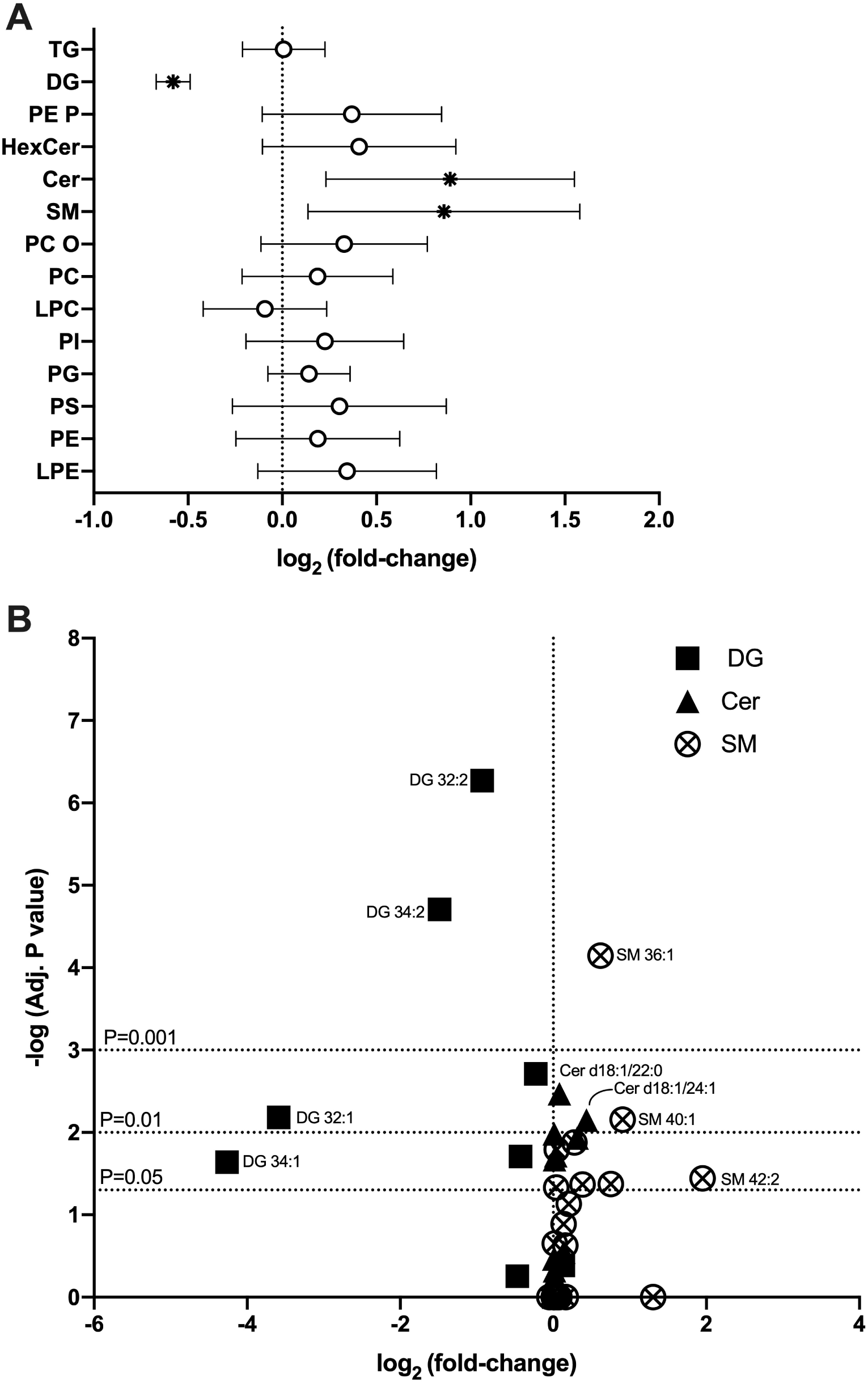
The effect of miR-221-3p on adipocyte lipid composition. A) Total lipid classes in miR-221-3p mimic transfected mature adipocytes (n=4). Significantly altered classes are marked with an asterisk. B) Individual affected species in the significantly affected classes. The most affected species are identified. The three horizontal lines illustrate P values. The vertical dotted line indicates non-targeting siRNA (NT) in both panels. TG, triglycerides; DG, diglycerides; LPC, lyso-phosphatidylcholines; PE P, phosphatidylethanolamine plasmalogens; HexCer, hexosylceramides; Cer, ceramides; SM, sphingomyelins; PC O, phosphatidylcholine ethers; PC, phosphatidylcholines; PI, phosphatidylinositols; PG, phosphatidylglycerols; PS, phosphatidylserines; PE, phosphatidylethanolamines; LPE, lyso-phosphatidylethanolamines.

### Ceramide and sphingomyelin accumulation and diacylglycerol defect may be mediated by downregulation of SMPD1, ASAH1, MOGAT and ACLY

We were interested in the mechanisms that could have mediated the lipidome alterations observed in miR-221-3p transfected cells. Hence, we analyzed by qPCR mRNA levels of a number of enzymes mediating the synthesis of DG, Cer or SM, or the degradation of the latter two (Fig. 7A). The results were consistent with the idea that miR-221-3p overexpression dampened the degradation of Cer and SM, as suggested by suppression of acid ceramidases 1 and 2 (*ASAH1, 2*) and sphingomyelin phosphodiesterase 1 (*SMPD1*; Fig. 7A). The down-regulation of ASAH1 was also confirmed at the protein level (Fig. 7B). Considering mechanisms underlying the reduction of DG, *de novo* lipogenesis (DNL) assays (Fig. 7C) suggested that the reason lies most likely in reduced synthesis of DG either through DNL or through a salvage reaction from monoacylglycerol by MOGAT. The *MOGAT* mRNA was shown to be significantly suppressed (Fig. 7A) in cells overexpressing miR-221-3p. For the *de novo* pathway we analyzed ATP-citrate lyase (ACLY), an emerging cardiovascular therapy target, which catalyzes a reverse step of the Krebs cycle, converting citrate to acetyl-CoA, the precursor for DNL [44, 45]. A significant effect on ACLY mRNA was not observed (P=0.06; Fig 7A). However, western blot analysis showed a significant down-regulation of the ACLY protein (Fig. 7B). To conclude, these observations revealed candidate mediators of the impacts of miR-221-3p on adipocyte lipid metabolism, suppression of *SMPD1* and *ASAH1/2* providing a plausible explanation for the increase of Cer and SM, and that of *MOGAT* and *ACLY* for the reduction of DG. Of note, none of these four enzymes are predicted to be direct targets of miR-221-3p, but are most likely indirectly affected by the miRNA.

**Fig. 7.**
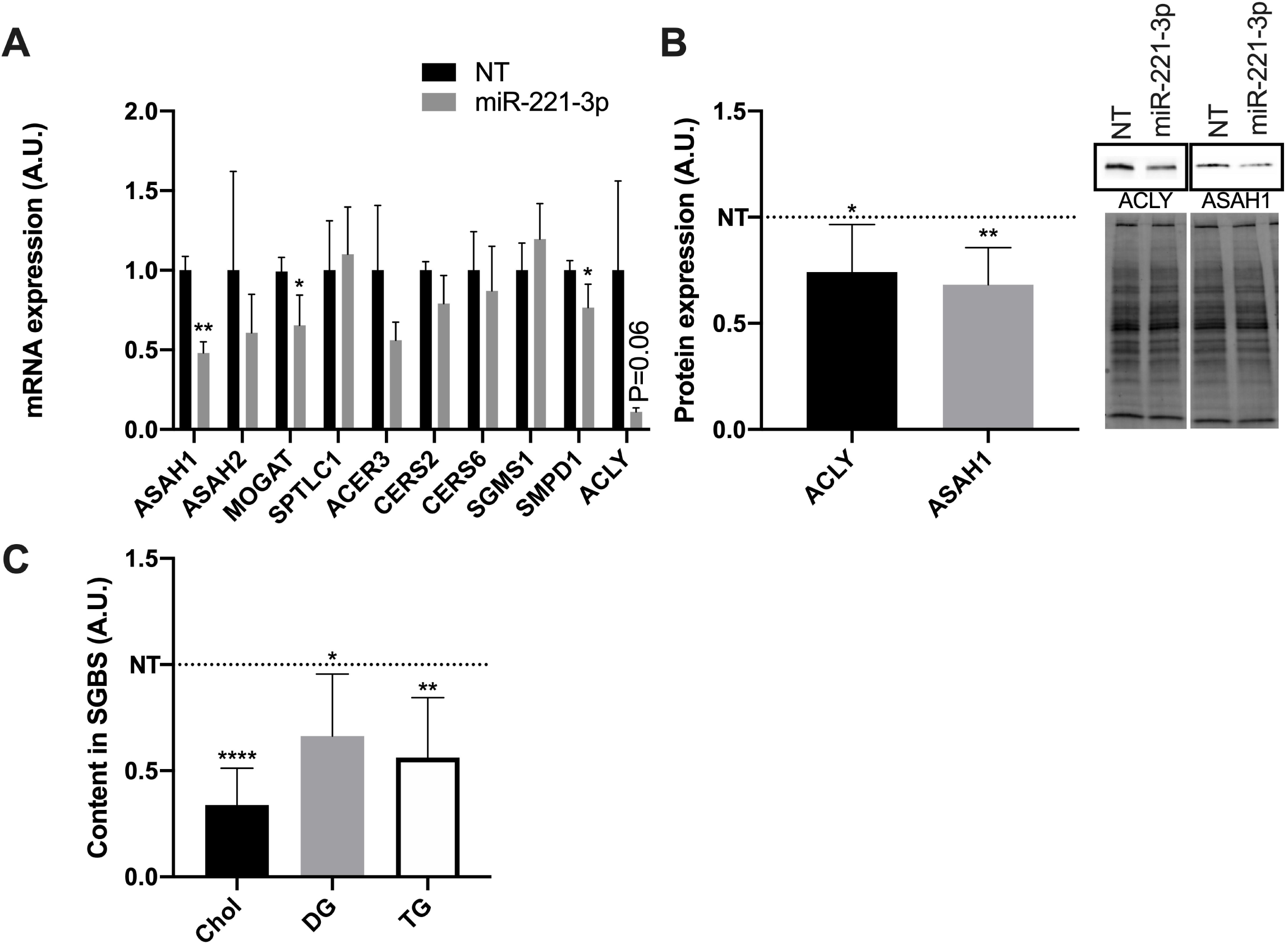
miR-221-3p overexpression inhibits ASAH1, MOGAT and ACLY expression and *de novo* lipogenesis. A) Expression of the indicated mRNAs in NT and miR-221-3p transfected adipocytes (n=3-6, two independent experiments). B) Protein expression of ASAH1 and ACLY in cells transfected with non-targeting siRNA (NT) or miR-221-3p mimic (miR-221-3p; n=9, three independent experiments). Representative western blots are shown on the right: immunodetected bands are shown at the top, and total protein at the bottom. C) *De novo* lipogenesis of cholesterol (Chol), DG and TG, as measured by [^3^H]acetic acid incorporation (n=12, two independent experiments). NT is represented by a dotted line. The data is represented as mean with SD. Statistical significance is designated as *\*P ≤ 0*.*05, **P < 0*.*01, ****P < 0*.*0001*.

### Mimicking tumor microenvironment enhances miR-221-3p expression

Elevation of miR-221-3p has been associated with several cancers [46–48] and it is thus considered an oncomiR. Since an altered phenotype characterized by impaired lipid storage is a characteristic of tumor-associated adipocytes [49, 50], we next studied the expression of miR-221-3p in the adipose tissue proximal to breast carcinoma (BC) in a total of 47 subjects undergoing mastectomy. Interestingly, the expression level of miR-221-3p increased along with the carcinoma grading, grade III displaying the highest expression of the miRNA (Fig. 8A). *AdipoQ*, which exerts anti-proliferative, anti-angiogenic and pro-apoptoptic effects [51], showed a significant negative correlation with the abundance of miR-221-3p (Fig. 8B). To study whether breast cancer cells could emit signals controlling the expression of miR-221-3p in adipocytes within the tumor microenvironment, we cultured SGBS adipocytes with MCF-7 BC (adenocarcinoma) cell line conditioned medium (CCM; Fig. 8C). Indeed, the CCM quite significantly enhanced miR-221-3p expression in the adipocytes (Fig. 8E). The increase of adipocyte miR-221-3p expression induced by CCM was abolished in the presence of actinomycin D (data not shown), suggesting that the miRNA elevation occurs via endogenous SGBS miRNA synthesis rather than through carry-over from the MCF-7 cells. qPCR analyses revealed that incubation of SGBS adipocytes with CCM significantly induced the mRNAs encoding the pro-inflammatory factors (visfatin; TIMP metallopeptidase inhibitor 1 - *Timp1*; *IL6* and *MCP1;* Fig. 8F). Furthermore, the adipogenic marker mRNAs *SCD1, GLUT4, DGAT1, ATGL*, and *AdipoQ* were significantly suppressed by the CCM treatment (Fig. 8G).

**Fig. 8.**
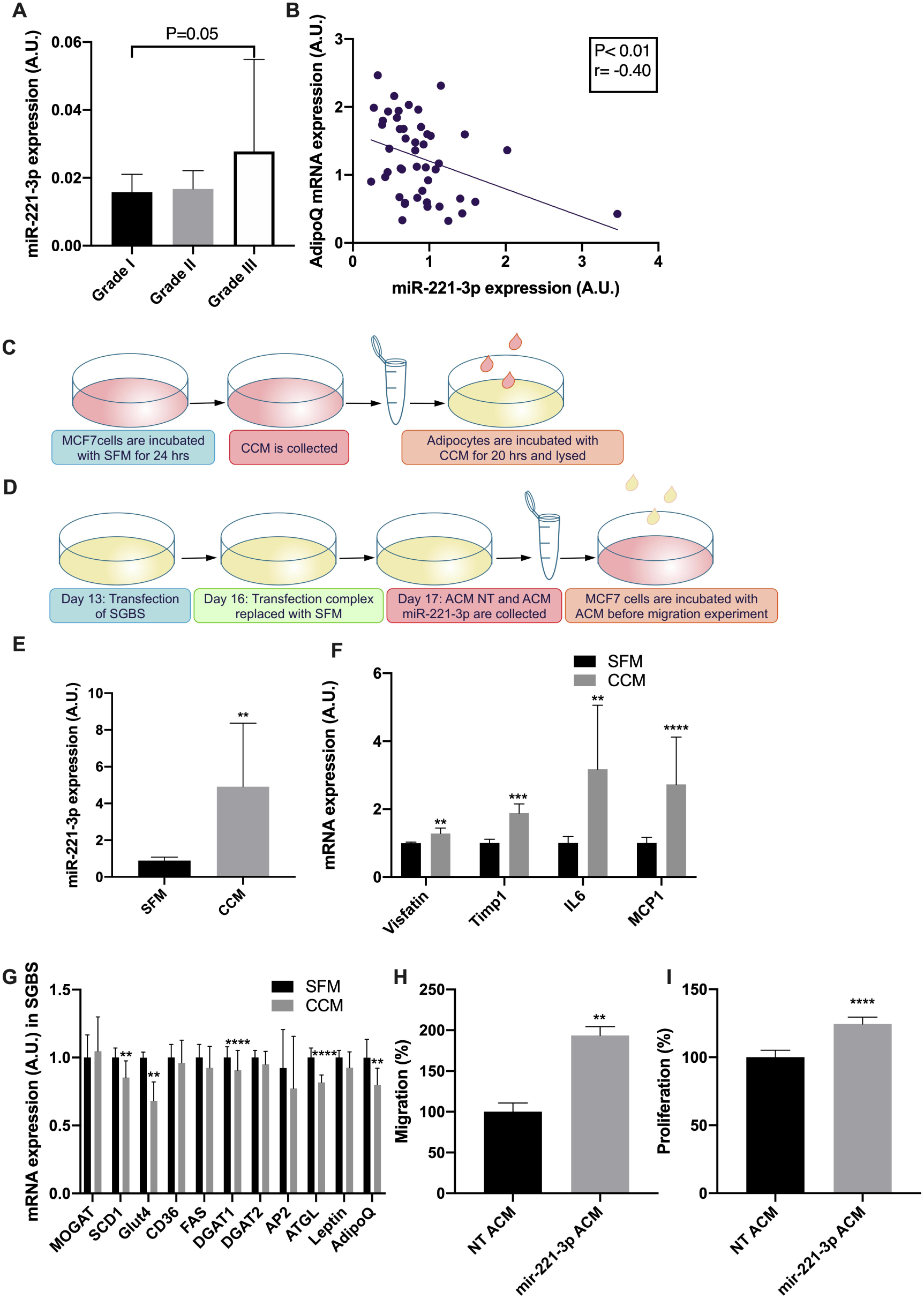
Role of miR-221-3p in the tumor microenvironment. A) miR-221-3p expression in tumor proximal adipose tissue of breast carcinoma patients (n=47; grade I, n=14; grade II, n=15; grade III, n=18). B) miR-221-3p negatively correlates with adiponectin (*AdipoQ*) mRNA in thesse breast adipose biopsies. P and r values are indicated in the panel (n=47). C-D) Outlines of *in vitro* medium transfer experiments. C) Cancer conditioned serum-free medium (SFM), designated CCM, was collected from MCF-7 breast cancer cells and incubated with SGBS adipocytes. D) Adipocyte conditioned SFM, designated ACM, was collected from mature SGBS transfected with non-targeting siRNA (NT) or miR-221-3p mimic (miR-221-3p), and incubated with MCF-7 cells. E) Expression of miR-221-3p (E), of inflammatory marker mRNAs (F) and of lipogenic marker mRNAs (G) in mature SGBS adipocytes treated with SFM or CCM (n=5-10, two to three independent experiments). Invasion (H) and proliferation (I) rate of MCF-7 cells treated with NT ACM or miR-221-3p ACM (n=6-9, three independent experiments). The data is represented as mean with SD. Statistical significance is designated as *\*\*P < 0*.*01,***P < 0*.*001, ****P < 0*.*0001*.

### Secretome of miR-221-3p overexpressing adipocytes increases invasion and proliferation of MCF-7 breast cancer cells

Since miR-221-3p expression was increased along with the progression of BC, we tested whether elevated expression of miR-221-3p in adipocytes might affect the phenotype of near-by BC cells. We therefore cultured MCF-7 cells with conditioned medium (ACM) from miR-221-3p or NT transfected SGBS adipocytes (Fig. 8D), and measured the migration and proliferation of the MCF-7 cells as endpoints related to cancer progression. Intriguingly, the secretome of miR-221-3p overexpressing adipocytes increased the migration rate of MCF-7 cells by 94% (Fig. 8H) and proliferation by 24% (Fig. 8I) when compared to ACM from adipocytes transfected with the NT control. This observation lends support to the idea that elevated expression of miR-221-3p in tumor-proximal, phenotypically altered adipose tissue may promote cancer progression.

## Discussion

Defects in adipocyte differentiation and lipid metabolism are associated with the pathogenesis of a number of common diseases [52–60]. miRNAs play a crucial role in maintaining adipocyte homeostasis, and they can also be used to identify novel molecular mechanisms underlying adipocyte function. miR-221 is one such miRNA, the expression of which declines during the course of adipocyte differentiation [24, 61, 62], which is up-regulated in the adipose tissue of obese human subjects [23], but is also associated with a number of cancers [46–48]. In this study, the effects of miR-221-3p on terminal adipocyte differentiation and the mechanisms involved were investigated. miR-221-3p or its anti-miR were transfected at an intermediate stage of adipocyte differentiation. The miRNA drastically inhibited the differentiation as evidenced by reduced lipid droplet accumulation and TG content, the anti-miR showing the reverse phenotypic effects (Fig. 1B-D). Genes involved in adipocyte differentiation and lipid synthesis were suppressed by miR-221-3p, which validates the differentiation defect observed (Fig. 2A). Previous studies have indicated that miR-221 dampens adipocyte differentiation when transfected at the initial stages of the process [24]. The impact of miR-221 overexpression on preadipocyte to adipocyte maturation was influenced by whether the cell origin was from lean or obese subjects: miR-221 decreased adipocyte differentiation only in preadipocytes from obese subjects while little effect was seen in preadipocytes of lean origin [63]. Importantly, these studies employed miR-221 expression at the initial and pre-initial stages of adipocyte differentiation. In the present study miR-221-3p was overexpressed at the intermediate stage of differentiation, thus targeting the terminal adipocyte differentiation relevant for the obesity-related dysfunction of adipocytes. Results of this and previous publications indicate that miR-221 is capable of inhibiting adipocyte differentiation at various stages of the differentiation cascade and might act through different target genes in a stage-dependent fashion. In obese and inflammatory conditions, adipocyte miR-221 expression is elevated [21, 23]. This elevation could contribute to an inhibition of healthy adipocyte expansion. Healthy adipocyte expansion and terminal differentiation are crucial to maintain AT insulin sensitivity and to limit the pathogenic hypertrophy of insulin resistant adipocytes [59, 64–66]. Moreover, elevated adipocyte miR-221 expression in tumor microenvironments, in which adipocytes are altered towards a more immature phenotype with reduced lipid storage capacity [49, 54], could promote tumor progression.

The mechanism by which miR-221-3p inhibits adipocyte differentiation and lipid droplet formation was analyzed. One of the predicted targets of miR-221-3p was 14-3-3γ, the expression of which increases during adipocyte differentiation [67]. In addition, 14-3-3γ was shown to be a direct target of miR-222, the target-selective seed sequence of which is identical to miR-221-3p [36]. Indeed, the expression of 14-3-3γ was markedly reduced in miR-221-3p overexpressing adipocytes (Fig. 3B), and 14-3-3γ knock-down suppressed the terminal differentiation of human adipocytes, indicating that 14-3-3γ may play an important role in the inhibition of adipocyte differentiation by miR-221-3p (Fig. 3C-E). Moreover, 14-3-3γ expression correlated positively with adipogenic and lipogenic gene expression in the subcutaneous (breast) adipose tissue of human subjects (Fig. 4B-E), supporting the *in vitro* findings. 14-3-3γ belongs to a family of related adaptor molecules, which affect the function and localization of a number of signaling components and transcription factors [68, 69]. 14-3-3 proteins are involved in a spectrum of cellular processes such as cell survival, motility, insulin signaling, GLUT4 translocation, and DNA damage and repair [70, 71]. Among the family members 14-3-3ζ was previously shown to accelerate visceral fat adipogenesis by affecting the key adipogenic transcription factors C/EBP-alpha and PPARγ in the initial phase of adipogenesis [35]. Moreover, 14-3-3ζ overexpression in mice led to a healthy expansion of adipocytes, which did not exhibit insulin resistance [35]. However, 14-3-3γ, in addition to being induced during adipocyte differentiation [67], is expressed at higher levels in obese adipose tissue compared to lean subjects [72, 73]. This observation is perplexing since miR-221 is also reported to be up-regulated in obesity [23]. Elevated expression of 14-3-3γ in obese conditions could represent a compensatory response to miR-221-3p overexpression, or it could reflect the abundance and high lipid content of adipocytes under conditions in which the AT is not yet dysfunctional. miR-221-3p inhibits adipocyte differentiation as shown during the present study, and miR-221-3p expression is induced by macrophage-mediated inflammation in adipocytes [21]. Altogether, our results may reflect for 14-3-3γ a scenario similar to that of 14-3-3ζ: The γ isoform could promote healthy AT expansion, and be up-regulated as a compensatory response when the AT becomes hypertrophic, inflamed and dysfunctional, a state increasing the abundance of miR-221-3p with an adverse effect on the adipocyte phenotype.

We next progressed to study the effect of miR-221-3p expression in mature adipocytes. Expression of the anti-inflammatory adipokine adiponectin was found to be decreased in the miR-221-3p overexpressing adipocytes. Adiponectin levels are reduced in insulin resistance, diabetes, and obesity [74, 75]. It is further shown to regulate adipogenesis and adipocyte lipid accumulation, thus preventing the accumulation of lipids at ectopic sites [76]. Circulating adiponectin is also shown to induce ceramidase activity by binding to adiponectin receptors (AdipoR) 1 and 2, resulting in the reduction of ceramides in cells expressing AdipoR1 and -R2 and improved whole-body insulin sensitivity [77]. Suppression of adiponectin expression in miR-221-3p overexpressing cells might thus have an impact on ceramide accumulation in adipocytes. In addition, miR-221 is connected to inflammation [12]. Similarly, we observed enhanced expression of inflammatory markers upon miR-221-3p overexpression which could promote sphingolipid accumulation [13]. Further studies were focused on identifying the lipid composition of miR-221-3p overexpressing mature adipocytes. No drastic reduction in TG content was observed in the miR-221-3p transfected cells, however, a reducing tendency was detectable at 72 hrs (Fig. 5D), concomitant with a suppression of lipogenic gene expression in these adipocytes (Fig. 5B). These observations indicated that a delipidation process might be underway. Due to technical limitations overexpression of miR-221-3p in mature adipocytes beyond 72 hrs was not possible.

Quantitative lipidome analysis was carried out to identify other disease-relevant lipid classes affected by miR-221-3p gain-of-function. The analysis revealed an increase of Cer and SM, while DG were significantly decreased (Fig. 6A). Gene expression analyses suggested that ceramide synthases were not altered, while the spingomyelin phosphodiesterase *SMPD1* and the acid ceramidases *ASAH1* and *-2* were downregulated. The accumulation of ceramides in miR-221-3p overexpressing mature adipocytes may thus be a result of ASAH1 downregulation, but also suppression of *AdipoQ* may contribute to this, as adiponectin stimulates ceramidase activity (see above paragraph). Increased ceramides, together with macrophage infiltration, are observed in subcutaneous AT of obese women with increased liver fat, suggesting that ceramides may amplify chronic AT inflammation and insulin resistance (37). Moreover, ceramide species including C14:0, C16:0, C16:1 and C18:1 are elevated in the white AT of obese and diabetic patients (11, 38-39). An increase of ceramides was also revealed upon lipidomic profiling of AT from subjects with cancer-related or primary lymphedema, reflecting a pro-inflammatory state of the AT (40). Our observations are consistent with the findings that ASAH1 overexpression in white AT improved whole-body insulin sensitivity and also alleviated hepatic steatosis. C16, C18, C24 and total ceramides were reduced in ASAH1 overexpressing adipose tissue [78]. Moreover, ceramides accumulate in pre-eclampsia due to a reduction of ASAH1 protein expression [79]. Even though ceramides have the capacity to induce apoptosis [80], we observed no such effect of miR-221-3p overexpression under the present experimental conditions (data not shown). To conclude, the increase of ceramides in the miR-221-3p overexpressing adipocytes is consistent with suppression of the adipocytes’ differentiated phenotype and insulin sensitivity, as well as a pro-inflammatory state.

The increase of sphingomyelins observed upon miR-221-3p overexpression could in principle be secondary to the increase of ceramides, since a conversion between sphingomyelins and ceramides occurs [81]. However, SM accumulation is often associated with an increase in DG, since the conversion of ceramides to SM by SM synthases occurs via transfer of phosphocholine from phosphatidylcholines (PC) to Cer, thus generating DG [82]. On the contrary, DG levels were diminished in miR-221-3p overexpressing adipocytes, and the SM phosphodiesteraae *SMPD1* was suppressed. This, together with the observed drop in DNL in these cells, indicates that the observed SM accumulation may not be a result of increased *de novo* synthesis but rather of a reduced turnover of the lipid. As another mechanism that could potentially reduce the cellular DG, we observed reduced *MOGAT* expression. This enzyme converts monoacylglycerol back to DG, thus replenishing the cellular DG pool [83]. The ceramide accumulation in miR-221-3p overexpressing adipocytes was attributed to a reduced expression of acid ceramidase (ASAH1) observed at both mRNA and protein levels. Further analysis also revealed that the miR-221-3p transfected adipocytes expressed reduced levels of ATP citrate lyase (ACLY) (Fig. 7), an enzyme that catalyzes the conversion of citrate to acetyl CoA, the precursor for DNL [84]. ACLY deficiency in primary adipocytes was shown to be associated with insulin resistance [85]. Moreover, ACLY was shown to activate the carbohydrate response element binding protein (ChREBP) and to regulate both DNL and glucose metabolism in brown adipocytes [86].

Delipidation and reduced adipogenesis of cancer-associated adipocytes are observed in many malignancies, including breast cancer [49, 54, 87]. Since miR-221-3p reduces adipocyte differentiation, the expression of miR-221-3p was analyzed in adipose tissue proximal to BC. miR-221-3p expression was elevated in the breast adipose tissue from patients with grade III invasive BC. AT adiponectin expression was reduced in the invasive BC and correlated negatively with miR-221-3p, indicating a defect in adipogenesis near the tumor. Plausibly, miR-221-3p could thus play a role in reducing adipogenesis in the vicinity of breast tumors and promote a delipidated adipocyte phenotype. Reduced adipogenesis might pave the way for tumor cells to migrate both by partitioning nutrients as well as by reducing spatial constraints [87, 88]. Moreover, modulation of adipokine expression in the tumor microenvironment could facilitate migration and proliferation of the cancer cells [89–91]. Interestingly, conditioned medium (ACM) obtained from miR-221-3p overexpressing adipocytes increased the invasion and proliferation of MCF-7 cells (Fig. 8H, I). Altered adipokine expression or enrichment of nutrients in the ACM might contribute to the enhanced BC cell invasion and proliferation. On the other hand, conditioned medium of MCF-7 cells enhanced the expression of miR-221-3p and pro-inflammatory factors, while reducing adipogenic marker mRNAs, in SGBS adipocytes, suggesting a two-way communication between the cancer cells and the near-by adipocytes, which could promote tumor progression.

In conclusion, our results demonstrate a pivotal role of miR-221-3p in the terminal differentiation of human white adipocytes and their lipid composition. Elevated expression of miR-221-3p impairs adipocyte lipid storage and differentiation, as well as modifies their content of key signaling lipids, Cer and DG. These alterations are relevant for metabolic diseases, but may also facilitate cancer progression.

## Ethics approval

This study was approved by the Ethics Committee of Helsinki University Central Hospital.

## Consent to participate

A written informed consent was acquired from all study subjects.

## Consent for publication

A written informed consent was acquired from all study subjects.

## Availability of data and material

Available from the authors upon request.

## Code availability

Not applicable.

## Acknowledgments

Riikka Kosonen, Anne Ahmanheimo, and Eeva Jääskeläinen are thanked for expert technical assistance.

